# Comparison of read mapping and variant calling tools for the analysis of plant NGS data

**DOI:** 10.1101/2020.03.10.986059

**Authors:** Hanna Marie Schilbert, Andreas Rempel, Boas Pucker

## Abstract

High-throughput sequencing technologies have rapidly developed during the past years and became an essential tool in plant sciences. However, the analysis of genomic data remains challenging and relies mostly on the performance of automatic pipelines. Frequently applied pipelines involve the alignment of sequence reads against a reference sequence and the identification of sequence variants. Since most benchmarking studies of bioinformatics tools for this purpose have been conducted on human datasets, there is a lack of benchmarking studies in plant sciences. In this study, we evaluated the performance of 50 different variant calling pipelines, including five read mappers and ten variant callers, on six real plant datasets of the model organism *Arabidopsis thaliana*. Sets of variants were evaluated based on various parameters including sensitivity and specificity. We found that all investigated tools are suitable for analysis of NGS data in plant research. When looking at different performance metrices, BWA-MEM and Novoalign were the best mappers and GATK returned the best results in the variant calling step.

## 1. Introduction

As the basis of biological properties and heredity, the genome of a species is a valuable resource for numerous studies. However, there are subtle differences between individuals of the same species, which are of academic and economic interest as these determine phenotypic differences. Dropping sequencing costs boosted high-throughput sequencing projects, thus facilitating the analysis of this genetic diversity. Variations within the *A. thaliana* population were studied in the 1001 genomes project [1]. As the number of high-quality reference genome sequences rises continuously, the number of re-sequencing projects increases as well [2]. There are pan-genome projects for various species focusing on the genome evolution [3–5] and mapping-by-sequencing projects which focus on agronomically important traits of crops [6–9].

An accurate and comprehensive identification of sequence variants between a sample and the reference sequence is the major challenge in many re-sequencing projects [10]. The large amount and diverse nature of NGS-data types (as reviewed in [11]), the diversity of bioinformatics algorithms, and the quality of the reference genome sequence render the choice of the best approach challenging.

Variant calling pipelines often start with (I) the preprocessing of sequence reads, followed by (II) the alignment (mapping) of these reads to a reference sequence. Finally, the (III) identification (calling) of sequence variants is performed based on alignments. Each of these three steps can be carried out by various alternative programs using different algorithms, which influence the accuracy and sensitivity of the resulting variant set.

First, read processing can be required if the read quality is at least partially low. Some downstream tools require that sequence reads come with quality scores in a certain system, namely phred33 or phred64. The conversion between different systems is allowed by some read processing tools. Popular read processing tools are FastQC [12], htSeqTools [13], NGSQC [14], SAMStat [15], and Trimmomatic [16].

As the read mapping determines the quality of the alignment, it is arguably the most important step [10]. Sequence reads are aligned to a suitable, but not necessarily the best place in the genome sequence. Often, there is a trade-off between mapping speed and the quality of the resulting alignment [17]. Numerous mappers are available, which utilize different algorithms and criteria to generate alignments [18,10]. Consequently, the choice of tool and parameters can have a large influence on the outcome of the mapping [19,20]. Reads originating from PCR duplicates should be remove from the mapping prior to the variant calling to improve reliability of the results [20]. Moreover, the quality of the reference genome sequence plays an important role for the performance of the mapper. Particular challenges are low-complexity sequences, repetitive regions, collapsed copies of sequences, contaminations, or gaps in the reference genome sequence [10]. Frequently applied read mappers are Bowtie2 [21], BWA-MEM [22], CLC Genomics Workbench (Quiagen), GEM3 [23], Novoalign (http://novocraft.com/), and SOAP2 [24]. While most of these tools are freely available for academic use as command line versions, CLC Genomics Workbench is a proprietary software suit for genomics with a graphical user interface. Detailed characteristics and algorithms of each mapper have been described elsewhere [25,26,20,18].

Finally, genomic variants like single nucleotide variants (SNVs) or small insertions/deletions (InDels) can be inferred by variant callers based on sequence read alignment. Popular variant callers like SAMtools/BCFtools [27], CLC Genomics Workbench (Quiagen), FreeBayes [28], GATK [29–32], LoFreq [33], SNVer [34], VarDict [35], and VarScan [36] use a variety of different approaches to call variants. Consequently, resulting variant sets differ depending on the employed methods (e.g. Bayesian), which come with strengths and weaknesses concerning the identification of specific variant types [10,37]. Several factors that contribute to high accuracy of variant callings are: (I) a high coverage of the variant position resulting in support for SNVs by several overlapping reads [38], (II) a careful study design [20], (III) joint variant calling for multiple samples to allow mutual support of genotypes [39].

The initial set of putative sequence variants is usually filtered to remove unreliable variant calls. Possible reasons for the identification of these variants in the first place are incorrect alignments, sequencing errors, or low-quality scores [10]. Read depth, mapping quality, and bias in the alignment to both strands are central criteria used in the filtering step. While this filter step aims to reduce the number of false positives, it simultaneously increases the number of false negatives. The best trade-off between sensitivity and specificity depends on the purpose of the respective study.

Many underlying algorithms of variant calling pipelines were developed for the analysis of variants in the human genome, e.g. to investigate genetic disorders or to study tumor samples [20,40– 43]. Although the applications in biomedical research and plant sciences differ substantially, plant scientists have largely followed benchmarking studies derived from research on human samples assuming similar performances. Moreover, many plant genomes possess unique challenges for variant calling, namely high amounts of repetitive sequences [44], large structural variations [45], broad range of heterozygosity and polyploidy [46]. Therefore, the diversity of plant genomes reveals the necessity of a benchmarking study using plant data sets. However, no comprehensive benchmarking study of read mapping and variant calling tools for plant genome sequences is described in the literature. Due to substantial differences in the nucleotide composition, a dedicated benchmarking on plant genome sequences is advised. A recent study compared the performance of BWA-MEM [22], SOAP2 [24], and Bowtie2 [21] with the two variant callers GATK [29–32] and SAMtools/BCFtools [27] on simulated and real tomato datasets [47]. To expand the sparse knowledge about the performance of other read mapping and variant calling tools on plant data, we set out to perform a systematic comparison. Due to the availability of excellent genomic resources, we selected the well-established plant model organism *Arabidopsis thaliana* for our study. While *A. thaliana* is not perfectly representing all plants, the genome shows the characteristic high proportion of AT. Despite limitations in heterochromatic and centromeric regions [48], many plant repeats are resolved in the high quality genome sequence of *A. thaliana*. Our study evaluated the performance of 50 variant calling pipelines, testing combinations of five read mappers (Bowtie2, BWA-MEM, CLC-mapper, GEM3, Novoalign) and eight different variant callers (SAMtools/BCFtools, CLC-caller, FreeBayes, GATK v3.8/v4.0/v4.1, LoFreq, SNVer, VarDict, VarScan) that are frequently applied in re-sequencing studies. Many combinations perform almost equally well on numerous datasets of the plant model organism *A. thaliana*. Illumina sequence reads were used for the detection of variants and provide the foundation for the comparison of these pipelines. Independent PacBio long reads were harnessed for the validation of identified variants based on an orthogonal sequencing technology.

## 2. Results

### 2.1. General stats about mapping of reads

Six Illumina paired-end sequence read datasets [49,50] from *A. thaliana* Nd-1 and one control sample of Col-0 [49] were processed using all combinations of five read mapping and eight different variant calling tools (including three different versions of one tool) to evaluate the mapping percentage as well as precision, sensitivity, and specificity of each variant calling pipeline. Due to these combinations (7×5×10), 350 variant calling sets were generated in this study. Overall, the sequence read quality of the processed datasets was high ranging from an average Phred score of 35 to 38 (Table S1).

We observed only minor differences between the different sequence read datasets with respect to percentage of properly aligned read pairs (Figure S1). In general, a higher proportion of the 2×300 nt paired-end reads was mapped ranging from 94.8% to 99.5%, while the 2×250 nt and the 2×100 nt paired-end reads resulted in mapping proportions ranging from 92.7% to 99.6% and 90.0% to 99.1%, respectively.

The comparison of mapping performance revealed that GEM3 had the highest average percentage of aligned read pairs (99.4%), followed by Novoalign (98.8%), Bowtie2 (98.5%), BWA-MEM (98.1%), and the read mapping function within CLC Genomics Workbench (CLC-mapper) (92.9%) (Figure 1).

**Figure 1.**
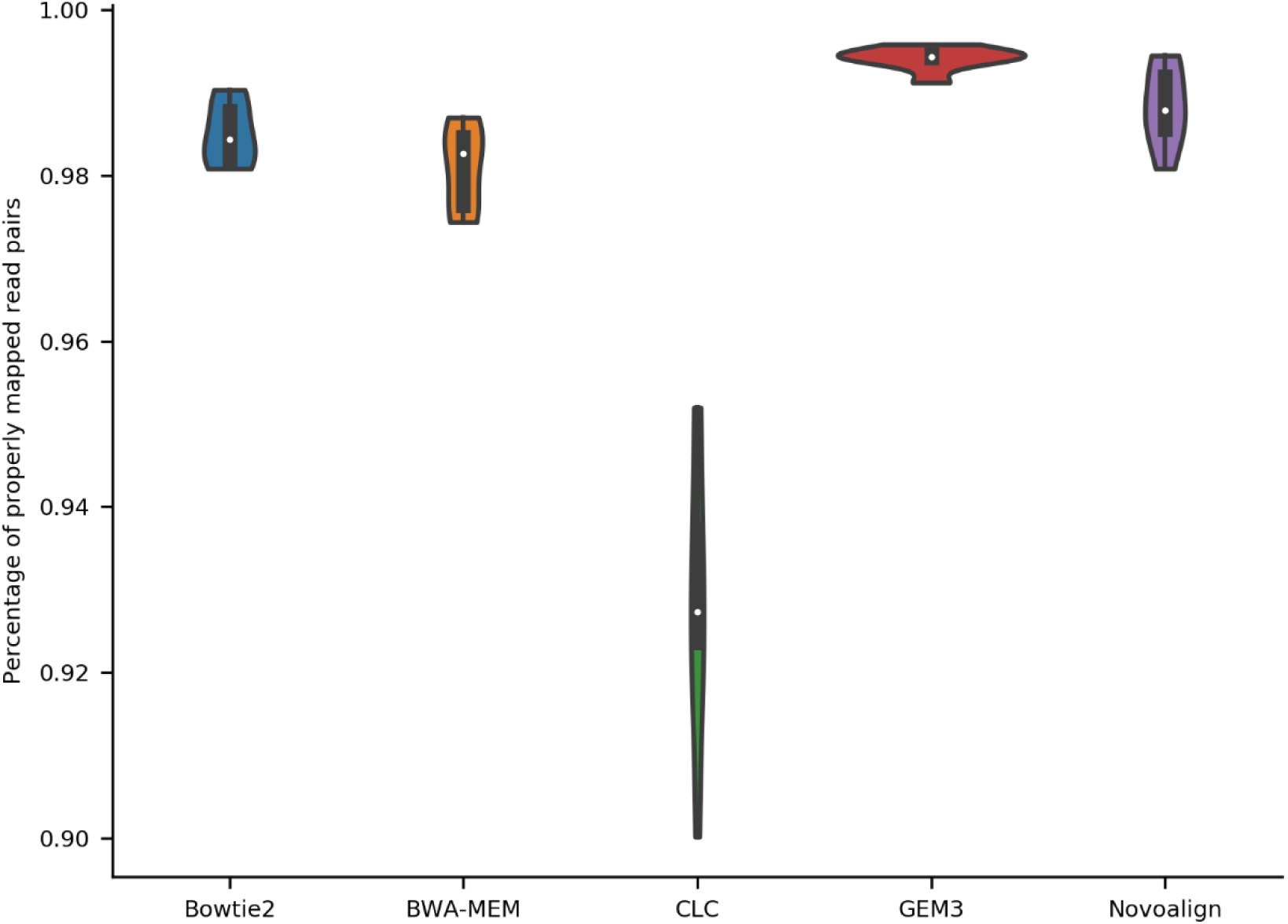
Ratio of mapped sequence read pairs per mapper. Sequence reads of six *A. thaliana* Nd-1 datasets were mapped to the Col-0 reference genome sequence TAIR10. The average ratio of aligned read pairs was calculated for Bowtie2, BWA-MEM, the mapping function in CLC Genomics Workbench (CLC), GEM3, and Novoalign based on all six datasets through the flagstats function of SAMtools. The width of the violin plots is proportional to the density of the data points. The boxplots inside the violin plots indicate quantiles and the white dot indicates the median.

### 2.2. Initial variant calling results & validation results

The initial variant calling revealed between 32,939 (Bowtie2 / CLC-caller) and 1,009,163 (BWA-MEM / VarDict) unfiltered SNVs, while the number of unfiltered InDels ranged from 2,559 (BWA-MEM / VarScan) to 240,879 (GEM3 / VarDict) (Table S2). Based on the three variant callers, which were able to call and classify MNVs (CLC-caller, VarDict, and FreeBayes), MNVs ranged from 1,394 (Bowtie2 / CLC-caller) to 168,100 (CLC-mapper / FreeBayes) (Table S2).

The quality of a variant call set is determined by the transition/transversion ratio, as a worse variant call set tends to have a lower transition/transversion ratio [49]. While most variant callers showed a similar transition/transversion ratio with a median ranging from 1.256 (LoFreq) to 1.288 (VarDict), SNVer revealed a lower median of 1.2 and especially FreeBayes performed worst, showing a median of 1.15 (Figure 2). In addition, FreeBayes revealed the greatest variation ranging from 0.92 to 1.31.

**Figure 2.**
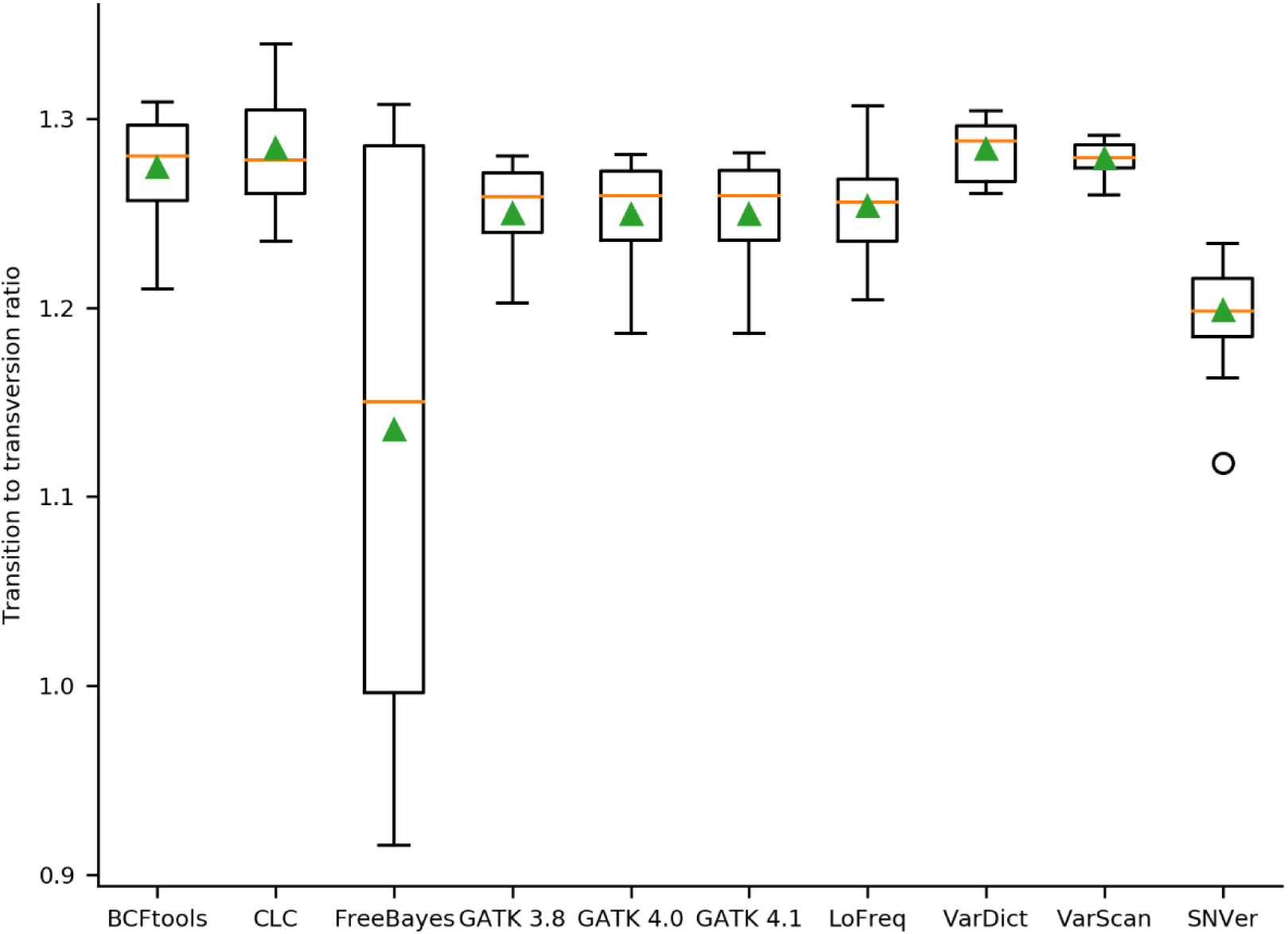
Ratio of transitions/transversions in the variant call sets per variant caller. Evaluation of call set quality was harnessed by analyzing the transition/transversion ratio. The orange line represents the median, the green triangle represents the mean.

In order to analyze whether a variant caller identifies relatively more SNVs than InDels, the ratio of SNVs to SNVs and InDels was calculated per variant caller (Figure 3). BCFtools identified the highest proportion of SNVs (median = 0.90), while VarDict and GATK 4.1 called the lowest proportion of SNVs (median = 0.824). Moreover, all GATK versions performed similar and revealed low variance when compared to the other variant callers. Interestingly, BWA-MEM / VarScan using the SRR3340910 dataset yielded the highest SNVs/InDels ratio with almost 1 (0.996).

**Figure 3.**
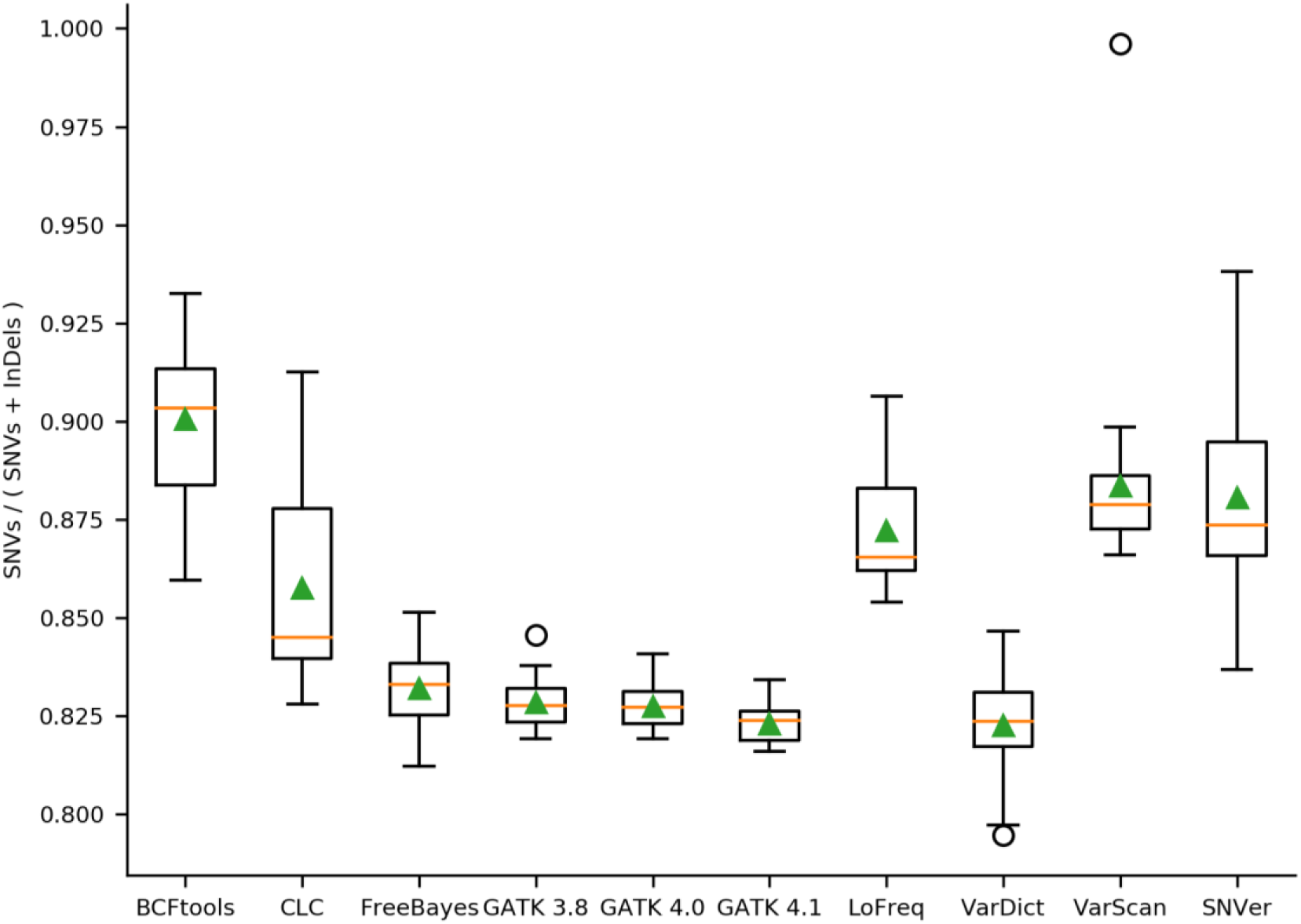
Proportion of SNVs to all variants per variant caller. Performance of each variant caller was assessed based on 30 mappings of *A. thaliana* Nd-1 reads against the Col-0 reference genome sequence TAIR10. Evaluation of the proportion of SNVs to all variants in the results of each applied variant caller was analyzed. MNPs were excluded because not all variant callers identified MNPs. The orange line represents the median, the green triangle the mean.

To infer whether a variant caller is more prone to call small or large InDels, the distribution of InDel lengths was inspected (Figure 4). Especially VarDict called very large insertions (up to 981 bp) and very large deletions (up to 998 bp), which are likely to be artifacts since they are filtered out in the corresponding validated call set (Figure S2). VarScan (134 to −93), SNVer (134 to −95), CLC-caller (156 to −95), LoFreq (168 to −109), and BCFtools (216 to −108) showed a narrower range of InDel lengths.

**Figure 4.**
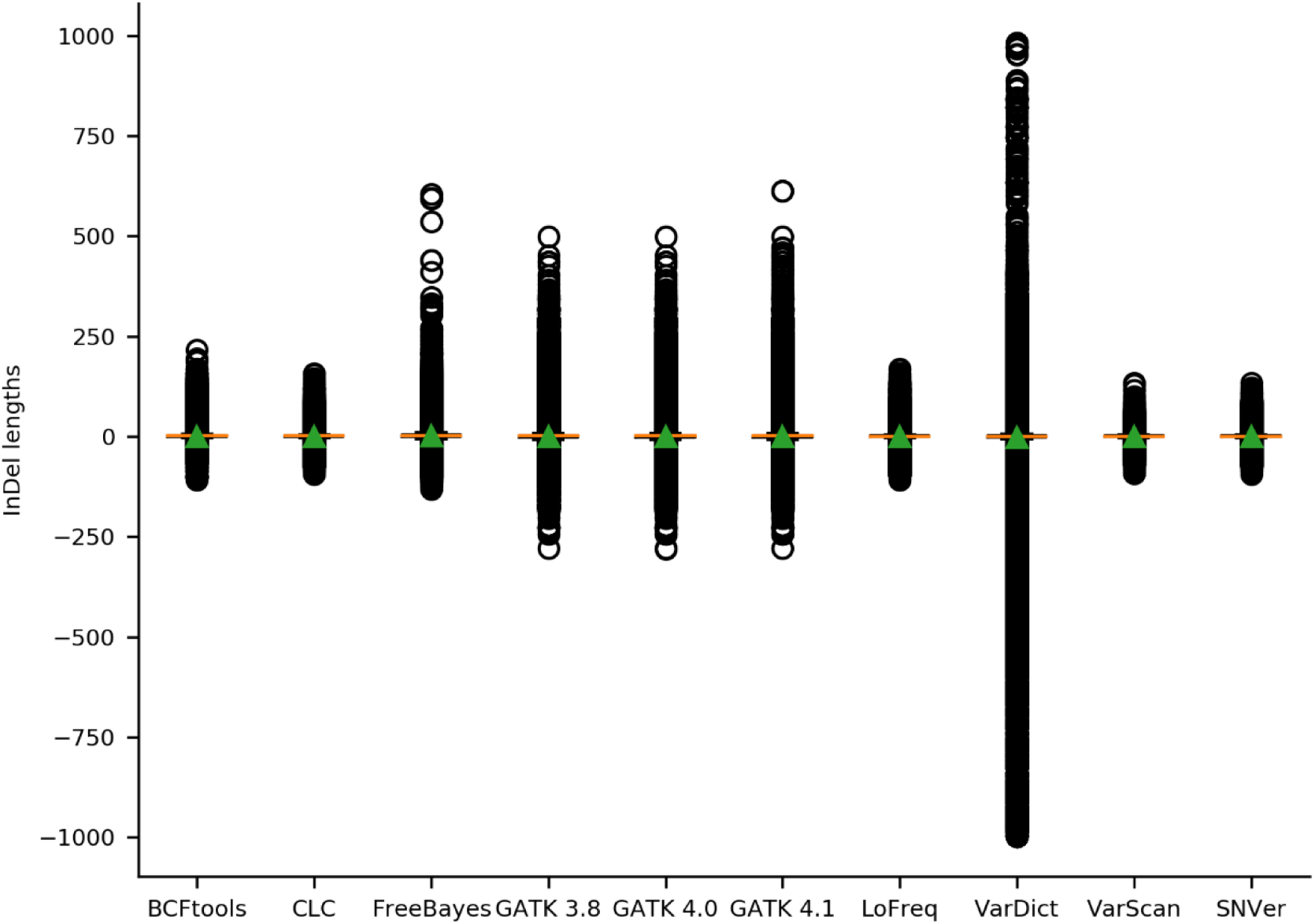
Distribution of InDel lengths per variant caller. Performance of variant callers was assessed based on 30 mappings of *A. thaliana* Nd-1 reads against the Col-0 reference genome sequence TAIR10. The length distribution of all InDels identified by each applied variant caller was analyzed. The orange line represents the median, the green triangle represents the mean.

A gold standard comprising variants which have been validated through orthogonal data was used for benchmarking (see methods for details). In order to compare the performance of different variant calling pipelines, we calculated the sensitivity, specificity, accuracy, precision, and F1 score (Table 1, Table S2). GATK revealed the highest accuracy in combination with most mappers. The only exception is the combination of GEM3 and VarScan, which performed better than any GATK version (Figure 5). GATK worked best on alignments produced by BWA-MEM and Novoalign. All three evaluated GATK versions (v3.8, v4.0, and v4.1) showed an almost identical performance. In general, Novoalign reached the best (median) results with respect to accuracy. The only exceptions are CLC-caller and VarScan based on alignments produced by CLC-mapper and GEM3, respectively. While Bowtie2 was identified to yield high specificity with most variant callers, it showed a low accuracy with most variant callers except for FreeBayes and VarDict.

**Table 1.** Performance statistics of variant calling pipelines. For each variant calling pipeline the statistics to infer performance are listed. sen = sensitivity, spe = specificity, pre = precision, acc = accuracy, F1 = F1 score.

|  | BCFtools | CLC | FreeBayes | GATK 3.8 | GATK 4.0 | GATK 4.1 | LoFreq | SNVer | VarDict | VarScan |
| --- | --- | --- | --- | --- | --- | --- | --- | --- | --- | --- |
| **Bowtie2** | 26.701 sen | 7.979 sen | 43.729 sen | 33.836 sen | 33.819 sen | 33.869 sen | 12.282 sen | 21.844 sen | 37.343 sen | 13.910 sen |
|  | 99.928 spe | 99.987 spe | 99.566 spe | 99.937 spe | 99.937 spe | 99.936 spe | 99.985 spe | 99.969 spe | 99.884 spe | 99.979 spe |
|  | 81.800 pre | 88.244 pre | 53.735 pre | 87.286 pre | 87.342 pre | 87.035 pre | 91.607 pre | 90.107 pre | 79.803 pre | 89.530 pre |
|  | 99.030 acc | 98.828 acc | 98.939 acc | 99.104 acc | 99.104 acc | 99.104 acc | 98.878 acc | 98.983 acc | 99.120 acc | 98.894 acc |
|  | 0.403 F1 | 0.145 F1 | 0.47 F1 | 0.488 F1 | 0.488 F1 | 0.488 F1 | 0.214 F1 | 0.352 F1 | 0.512 F1 | 0.239 F1 |
| **BWA-MEM** | 43.186 sen | 34.992 sen | 43.053 sen | 47.130 sen | 47.099 sen | 47.136 sen | 38.874 sen | 41.435 sen | 40.302 sen | 37.506 sen |
|  | 99.806 spe | 99.908 spe | 99.548 spe | 99.842 spe | 99.842 spe | 99.839 spe | 99.892 spe | 99.868 spe | 99.852 spe | 99.907 spe |
|  | 73.182 pre | 81.892 pre | 51.289 pre | 79.129 pre | 79.083 pre | 78.786 pre | 81.122 pre | 79.521 pre | 76.686 pre | 83.535 pre |
|  | 99.120 acc | 99.099 acc | 98.916 acc | 99.179 acc | 99.179 acc | 99.178 acc | 99.122 acc | 99.133 acc | 99.124 acc | 99.120 acc |
|  | 0.543 F1 | 0.492 F1 | 0.459 F1 | 0.591 F1 | 0.59 F1 | 0.59 F1 | 0.528 F1 | 0.546 F1 | 0.531 F1 | 0.518 F1 |
| **CLC** | 47.682 sen | 36.382 sen | 41.564 sen | 49.237 sen | 49.357 sen | 49.413 sen | 41.730 sen | 45.352 sen | 39.508 sen | 41.349 sen |
|  | 99.695 spe | 99.851 spe | 99.517 spe | 99.817 spe | 99.815 spe | 99.812 spe | 99.821 spe | 99.759 spe | 99.857 spe | 99.847 spe |
|  | 65.903 pre | 73.878 pre | 48.854 pre | 77.341 pre | 77.183 pre | 76.901 pre | 73.611 pre | 70.493 pre | 76.995 pre | 76.971 pre |
|  | 99.064 acc | 99.062 acc | 98.882 acc | 99.175 acc | 99.175 acc | 99.174 acc | 99.066 acc | 99.060 acc | 99.112 acc | 99.100 acc |
|  | 0.549 F1 | 0.491 F1 | 0.441 F1 | 0.601 F1 | 0.602 | 0.602 F1 | 0.536 F1 | 0.549 F1 | 0.525 F1 | 0.539 F1 |
| **GEM3** | 35.488 sen | 32.826 sen | 44.104 sen | 40.895 sen | 40.890 sen | 40.963 sen | 33.867 sen | 34.710 sen | 39.131 sen | 41.502 sen |
|  | 99.887 spe | 99.932 spe | 99.523 spe | 99.902 spe | 99.902 spe | 99.900 spe | 99.941 spe | 99.925 spe | 99.847 spe | 99.904 spe |
|  | 79.533 pre | 85.457 pre | 51.481 pre | 84.189 pre | 84.221 pre | 83.863 pre | 87.657 pre | 85.573 pre | 75.648 pre | 84.315 pre |
|  | 99.092 acc | 99.095 acc | 98.922 acc | 99.157 acc | 99.157 acc | 99.156 acc | 99.107 acc | 99.097 acc | 99.100 acc | 99.169 acc |
|  | 0.49 F1 | 0.475 F1 | 0.464 F1 | 0.55 F1 | 0.551 F1 | 0.55 F1 | 0.489 F1 | 0.494 F1 | 0.519 F1 | 0.557 F1 |
| **Novoalign** | 44.160 sen | 33.858 sen | 42.986 sen | 48.599 sen | 48.575 sen | 48.620 sen | 38.892 sen | 41.436 sen | 39.863 sen | 38.170 sen |
|  | 99.820 spe | 99.922 spe | 99.610 spe | 99.846 spe | 99.845 spe | 99.842 spe | 99.905 spe | 99.885 spe | 99.860 spe | 99.922 spe |
|  | 75.260 pre | 84.222 pre | 54.799 pre | 80.019 pre | 79.951 pre | 79.643 pre | 83.510 pre | 81.968 pre | 77.296 pre | 85.862 pre |
|  | 99.147 acc | 99.097 acc | 98.966 acc | 99.202 acc | 99.202 acc | 99.200 acc | 99.137 acc | 99.149 acc | 99.127 acc | 99.144 acc |
|  | 0.556 F1 | 0.483 F1 | 0.472 F1 | 0.605 F1 | 0.604 F1 | 0.604 F1 | 0.532 F1 | 0.551 F1 | 0.529 F1 | 0.529 F1 |

**Figure 5.**
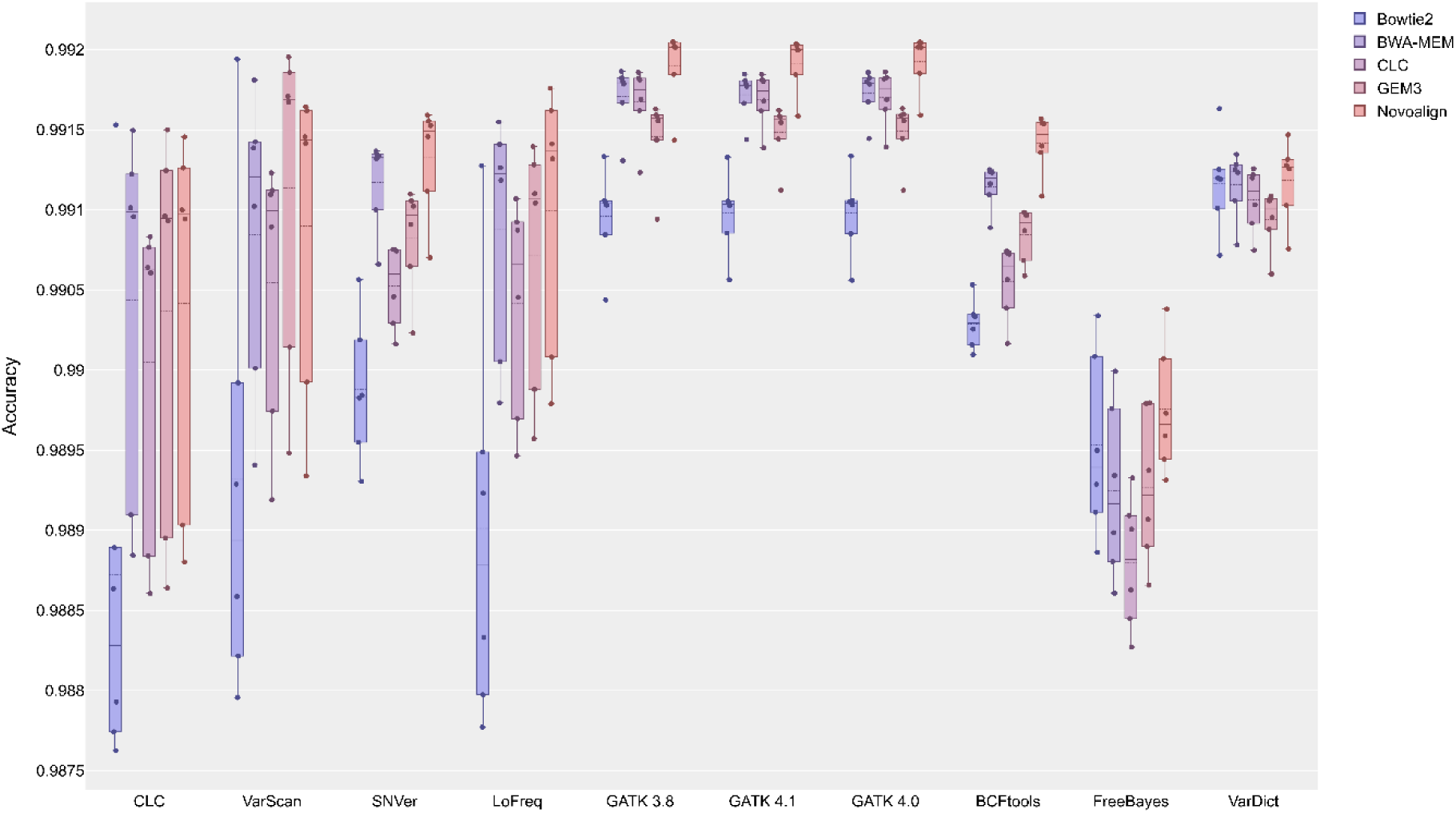
Accuracy of variant calling pipelines. The accuracy for each variant calling pipeline is shown with mean (dashed line) and median (straight line) calculated based on the results of the six analyzed datasets.

In general, the sensitivity of the variant caller pipelines ranged from 0.0219 (Bowtie2 / CLC-caller) to 0.6038 (BWA-MEM / VarDict) and the specificity from 0.99450 (CLC-mapper / FreeBayes) to 0.999961 (Bowtie2 / CLC-caller) (Figure S3, Figure S4). Moreover, we observed a negative correlation of −0.8 between specificity and sensitivity, indicating that a pipeline with a high sensitivity showed a low specificity and vice versa. Almost every variant caller, except for VarDict, showed the lowest specificity when used in combination with CLC-mapper, while in parallel these combinations had one of the highest sensitivities. VarDict showed the highest specificity, but lowest sensitivity with Bowtie2 and performed inferior to GEM3 in terms of specificity, while BWA-MEM reached the best results in sensitivity.

All tested GATK versions (v3.8, v4.0, and v4.1) showed a very high sensitivity and were only outperformed by specific VarDict samples, namely the 2×100 nt paired-end dataset independent of the choice of the mapper, which reached up to 0.6038 sensitivity (BWA-MEM / VarDict-SRR2919279). However, the specificity of GATK was inferior to some other variant callers. Only minor differences were observed between the three evaluated GATK versions regarding both sensitivity and specificity. The use of different mappers had a substantially higher impact than the applied GATK version.

Followed by GATK, FreeBayes showed a good performance in terms of sensitivity and robust results across all mappers, whereas the other variant callers showed a low performance in combination with Bowtie2. CLC-caller, VarScan, and LoFreq revealed a great variation with respect to sensitivity across all mappers, while GATK and especially VarDict displayed very low variance in their results. When focusing on median sensitivity, the following combinations showed the best results: CLC-mapper / CLC-caller, GEM3 / VarScan, CLC-mapper / SNVer, CLC-mapper / LoFreq, CLC-mapper / GATK v3.8, CLC-mapper / GATK v4.0, CLC-mapper / GATK v4.1, CLC-mapper / BCFtools, GEM3 / FreeBayes, and BWA-MEM / VarDict. However, in terms of median specificity all variant callers revealed the best results in combination with Bowtie2, except for FreeBayes, which worked best with Novoalign. Moreover, FreeBayes showed the lowest performance and largest variation across all mappers (Figure S3).

Finally, the harmonic mean of precision and sensitivity, namely the F1 score, was analyzed (Figure S5). Novoalign in combination with GATK revealed the best mean performance with respect to the F1 score. Again, different GATK versions showed almost identical performance (Table 1).

## 3. Discussion

The major challenge in many pangenome and re-sequencing projects is the accurate and comprehensive identification of sequence variants. Due to the high diversity and complexity of plant genomes and their differences to animal (e.g. human) genomes, variant callings in plant research differ substantially from those in human and biomedical research. Most human benchmarking studies focus on calling SNVs of certain tumor cells in a highly diverse cell set [20,41,42]. In contrast, plant studies usually use the whole cell set derived from one plant without genomic differences between cells. However, sequencing of pooled DNA from multiple plants aims at the identification of low frequency SNVs. Large amounts and different NGS data types (as reviewed in [11]), the diversity of bioinformatic algorithms, and the quality of the reference genome sequences render the choice of the best approach challenging. Hence, we performed a benchmarking study to provide comparable data showing the performance of combinations of frequently applied mappers and variant callers (variant calling pipelines) on plant datasets. A previous report [47] is extended by providing data about the performance of additional tools both for the mapping and variant calling step.

To allow for a consistent comparison of baseline performance, we used default parameters for all tools as these parameters are frequently chosen in plant science applications [4,5,9]. Sequence read datasets from different sequencing platforms, with different read lengths, and different sizes were processed to ensure a realistic benchmarking of tools. Since all evaluated tools can process a real plant dataset within a day, we refrained from assessing the computational costs of the analysis. There is usually a trade-off between quality of the results and computational costs. In our experience, many plant scientists select the workflow leading to the best results independent of computational costs [52].

The first step in a variant calling pipeline is the alignment (mapping) of reads to a reference sequence. While the mapping of 2×250 nt paired-reads resulted in a higher mapping percentage, the performance difference to 2×100 nt reads is only about 10%. As different sequencing platforms were used for the data generation, per base quality might contribute to this difference. It is expected that longer reads are aligned with higher specificity and hence improve the following variant calling.

The quality of the variant calling sets was assessed by the transition/transversion (ti/tv) ratio which was previously described as a quality indicator [51]. Overall, the quality of almost all analyzed call sets was similar. A previous benchmarking study with SAMtools and GATK reported similar ti/tv ratios for all pipelines [53]. A filtering step increased the ti/tv ratios, indicating that the filtering increased the quality of the call sets [53]. This observation is in line with our findings, which revealed an increased ti/tv for the filtered call sets reduced to variants present in the gold standard (Figure S6, Table S3). As FreeBayes showed a substantial increase in the quality through filtering, we recommend checking the transition/transversion ratio when applying FreeBayes. This effect might be dataset specific.

The choice of the variant caller is crucial for the number of called SNVs, MNVs, and InDels. For example, only CLC-caller, VarDict and FreeBayes were able to call MNVs, thus being more suitable for the identification of structural differences. Furthermore, variant caller results differ with respect to the ratio of SNVs to InDels, which should be considered depending on the specific requirements of the respective sequencing project. BCFtools called relatively more SNVs than InDels, while GATK revealed relatively more InDels. A unique property of VarDict was the detection of InDels up to almost 1 kb which exceeds the read length. Since the accurate identification of such large variants, which are longer than the average read length, is still a challenging task [54], many of these variants are likely false positives. Moreover, the reduced amount of large insertions in the validated call sets of VarDict supports this assumption.

Depending on the application, a pipeline with a high sensitivity or high specificity is desired. In terms of sensitivity, GATK in combination with CLC-mapper, Novoalign and BWA-MEM yielded the best and most consistent results across all evaluated datasets. These results are in line with a recent study showing that GATK often outperformed SAMtools in terms of sensitivity, precision, and called raw InDels [47]. Similar results had been shown in rice, tomato, and soybean [47] indicating that GATK is also suitable for various crop species with complex genomes. A high sensitivity is essential when a high number of true positives variations accelerates the power of the analyses, e.g. when looking for a detrimental variation between two samples. In this study, a pipeline comprising Bowtie2 and LoFreq resulted in the highest specificity and can thus be recommended. In contrast, a high specificity is desired in mapping-by-sequencing (MBS) projects, as a high proportion of true positives can keep the signal to noise ratio high. Combining both performance metrics by analyzing the accuracy, best results were obtained with Novoalign and GATK. The same pipeline yielded the best results regarding the harmonic mean of precision and sensitivity (F1 score). Differences observed between the three evaluated GATK versions (v3.8, v4.0, and v4.1) were negligible. However, functionalities and computational performance might differ between these versions.

In summary, this benchmarking study provides insights into the strengths and weaknesses of different variant calling pipelines when applied on plant NGS datasets. Although the performance of all evaluated tools will differ between samples depending on properties of the read datasets and the genome sequence, we hope that our findings serve as a helpful guide.

## 4. Materials and Methods

### Sequence read datasets

We used paired-end Illumina reads from two different *A. thaliana* accessions, namely Columbia-0 (Col-0) and Niederzenz-1 (Nd-1) (Table S1). The read quality was checked via FastQC [12] (Table S1). Trimmomatic [16] was applied for light trimming and quality conversion to a Phred score of 33 if applicable. These datasets cover different Illumina sequencing platforms including GAIIX, MiSeq, and HiSeq 1500. While two datasets are the paired-end proportions of mate pair sequencing libraries (SRR2919288 and SRR3340911), these samples are 2×250 nt paired-end libraries.

### Sequence read mapping

We chose five popular read mappers, namely Bowtie2 v2.3.4.3 [21], BWA-MEM v0.7.17 [22], CLC v11 Genomics Workbench (Quiagen), GEM3 v3.6 [23], and Novoalign v3.09.01 (http://novocraft.com/) for this analysis. While most of these mappers are freely available for academic use, CLC is a proprietary software suit for genomic analyses. Paired-end reads were mapped against the TAIR10 reference genome sequence [55]. The executed commands for each tool can be found in table S4. SAMtools v.1.8 [27] was deployed for sorting of the BAM files. Reads originating from PCR duplicates where removed via MarkDuplicates in Picard-bf40841 (https://broadinstitute.github.io/picard/). Read groups or InDel qualities were assigned as these are required by some tools for downstream processing. While the plastome and chondrome sequences were included during the mapping step, variant caller performance was only assessed for the five nucleome sequences.

### Variant calling

All mapping results were subjected to variant calling via CLC v11 Genomics Workbench (Quiagen), FreeBayes v1.2.0 [28], Genome Analysis Tool Kit v3.8/v4.0/v4.1 HaplotypeCaller (GATK-HC) [29–32], LoFreq v2.1.3.1 [33], SAMtools v1.9 [27] in combination with BCFtools v1.9 (alias BCFtools in the following), SNVer [34], VarDict [35], and VarScan [36]. We evaluated three different versions of GATK in order to analyze whether the applied version has a high impact on the variant calling pipeline performance. The executed commands for each tool can be found in table S4.

### Performance measure of variant calling pipelines

The overall workflow of our benchmarking study is presented in Figure 6. We applied a previously described pipeline to validate sequence variants against the Nd-1 de novo assembly based on PacBio reads (https://github.com/bpucker/variant_calling) [56], which is crucial in order to assess the performance of each variant calling pipeline. This Nd-1 genome sequence assembly is of high quality due to a high PacBio coverage of about 112-fold and additional polishing with about 120-fold coverage of accurate short reads [52]. A gold standard was generated from all validated variants by combining them into a single VCF file (https://docs.cebitec.uni-bielefeld.de/s/GG4CYJ7PcEwMFAF). Afterwards, the numbers of true positives, true negatives, false positives, and false negatives were calculated based on the gold standard and the initial VCF files for each variant calling pipeline and dataset. Next, performance statistics including sensitivity, specificity, precision, accuracy, and F1 score were calculated per combination of mapper, variant caller, and dataset (Table 1).

**Figure 6.**
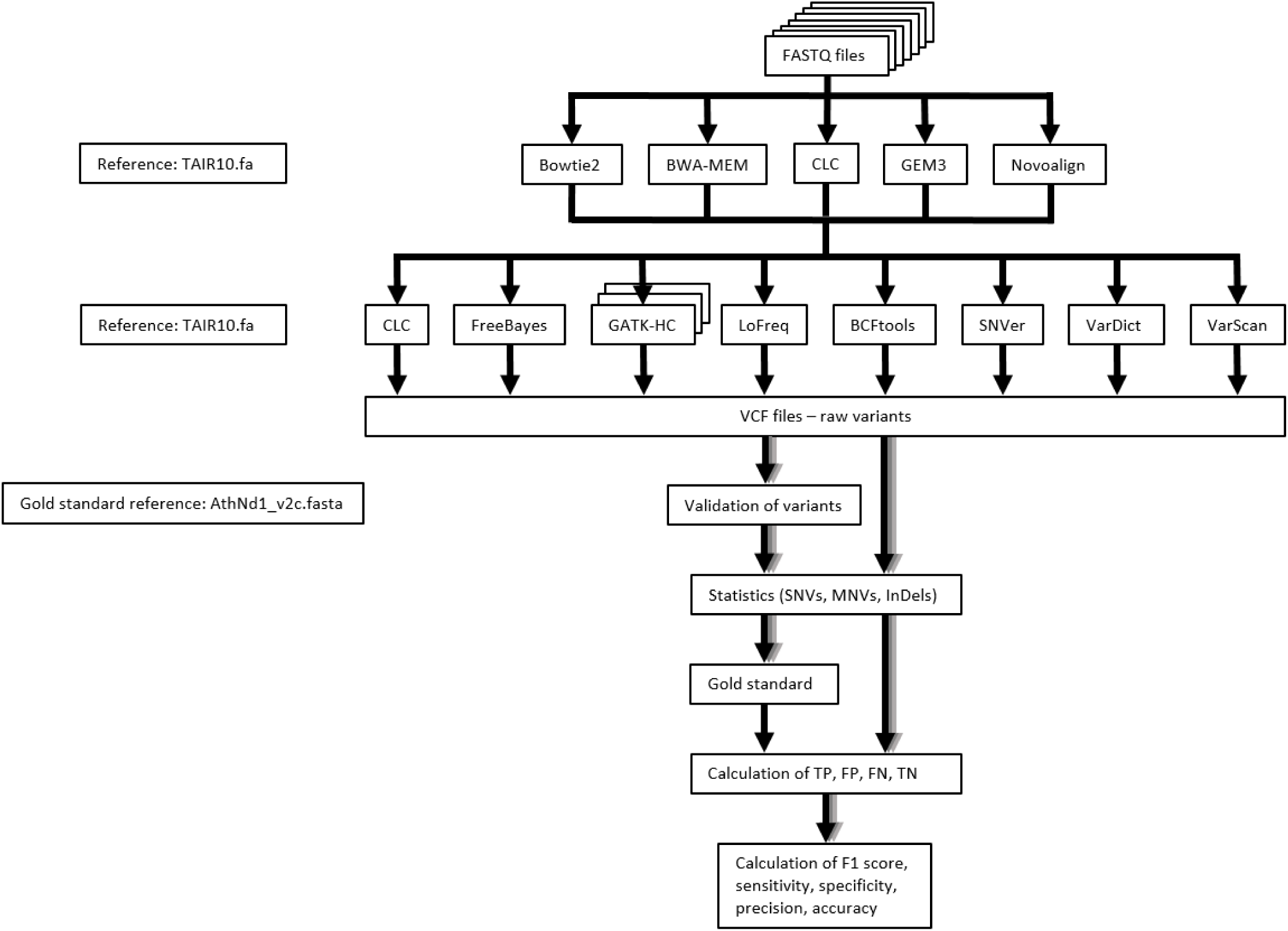
Workflow for the performance analysis of variant calling pipelines. First, reads within supplied FASTQ files were mapped against the TAIR10 *A. thaliana* reference genome sequence. Next, variants were called and saved in VCF files. All variants were subjected to a previously described validation process based on the Nd-1 genome sequence [56]. A gold standard was generated based on all validated variants. The initial variants called by each combination of mapper and variant caller were evaluated by analyzing whether they are present or absent in the gold standard. The numbers of SNVs, MNVs, and InDels were retrieved from the validated and from the initial VCF files (Table S2, Table S3). Next, true positives (TP), false positives (FP), false negatives (FN), and true negatives (TN) were calculated for SNVs, MNVs, and InDels identified by each combination of mapper and variant caller. Finally, performance statistics, such as F1 score, sensitivity, specificity, precision, and accuracy were calculated.

## Supplementary Materials

Table S1: Overview of used datasets with SRR identifiers. Table S2: Variant calling results of the initial VCF files. Table S3: Variant calling results of the validated VCF files. Table S4: Executed commands for each variant calling pipeline. Figure S1: Ratio of mapped read pairs per dataset. Figure S2: Distribution of InDel lengths in the validated call sets per variant caller. Figure S3: Specificity of variant calling pipelines. Figure S4: Sensitivity of variant calling pipelines. Figure S5: F1 score of variant calling pipelines. Figure S6: Ratio of transitions to transversions in the validated variant call sets per variant caller.

## Author Contributions

H.M.S., A.R., and B.P. conceived the project. H.M.S., A.R., and B.P. conducted data analysis. H.M.S. and B.P. wrote the manuscript. B.P. supervised the project. All authors have read and agreed to the published version of the manuscript.

## Funding

This research received no external funding.

### Acknowledgments

We thank the Bioinformatics Resource Facility support team of the CeBiTec for great technical support. We acknowledge support for the Article Processing Charge by the Open Access Publication Fund of Bielefeld University.

## Conflicts of Interest

The authors declare no conflict of interest.

## Notes

#### Summary of Updates

We added a more detailed description of the gold standard.

