## Supplementary figures and images for "Comparison of read mapping and variant calling tools for the analysis of plant NGS data"

### Figure S1

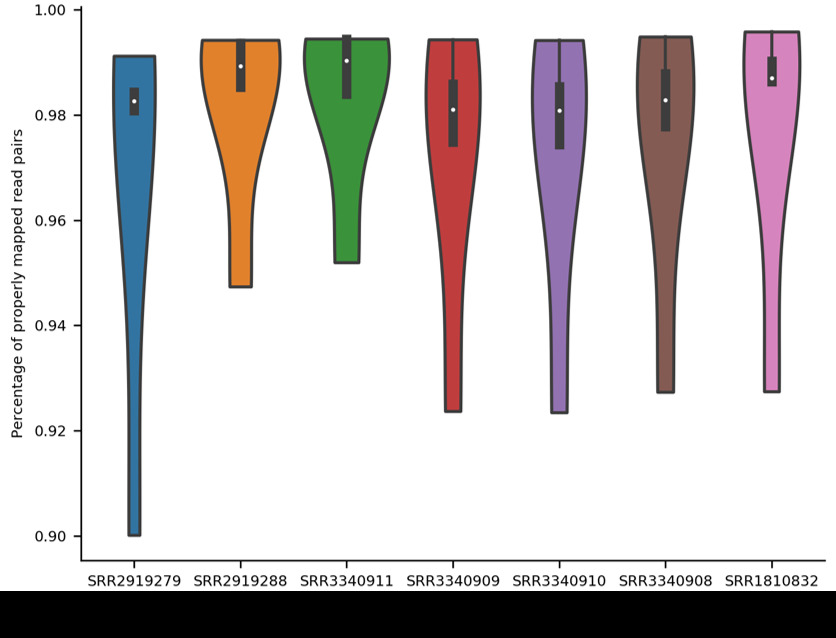

### Figure S2

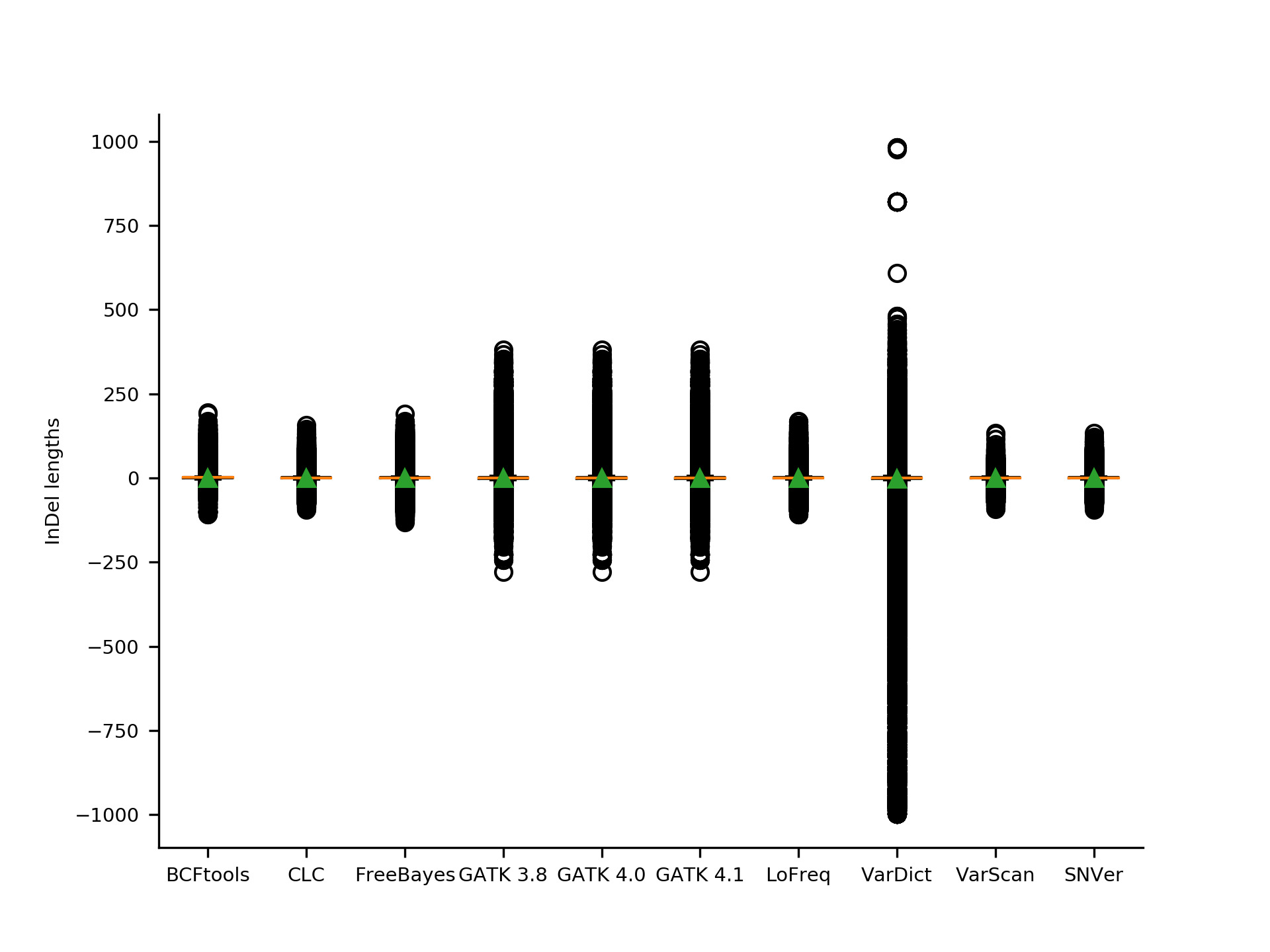

### Figure S3

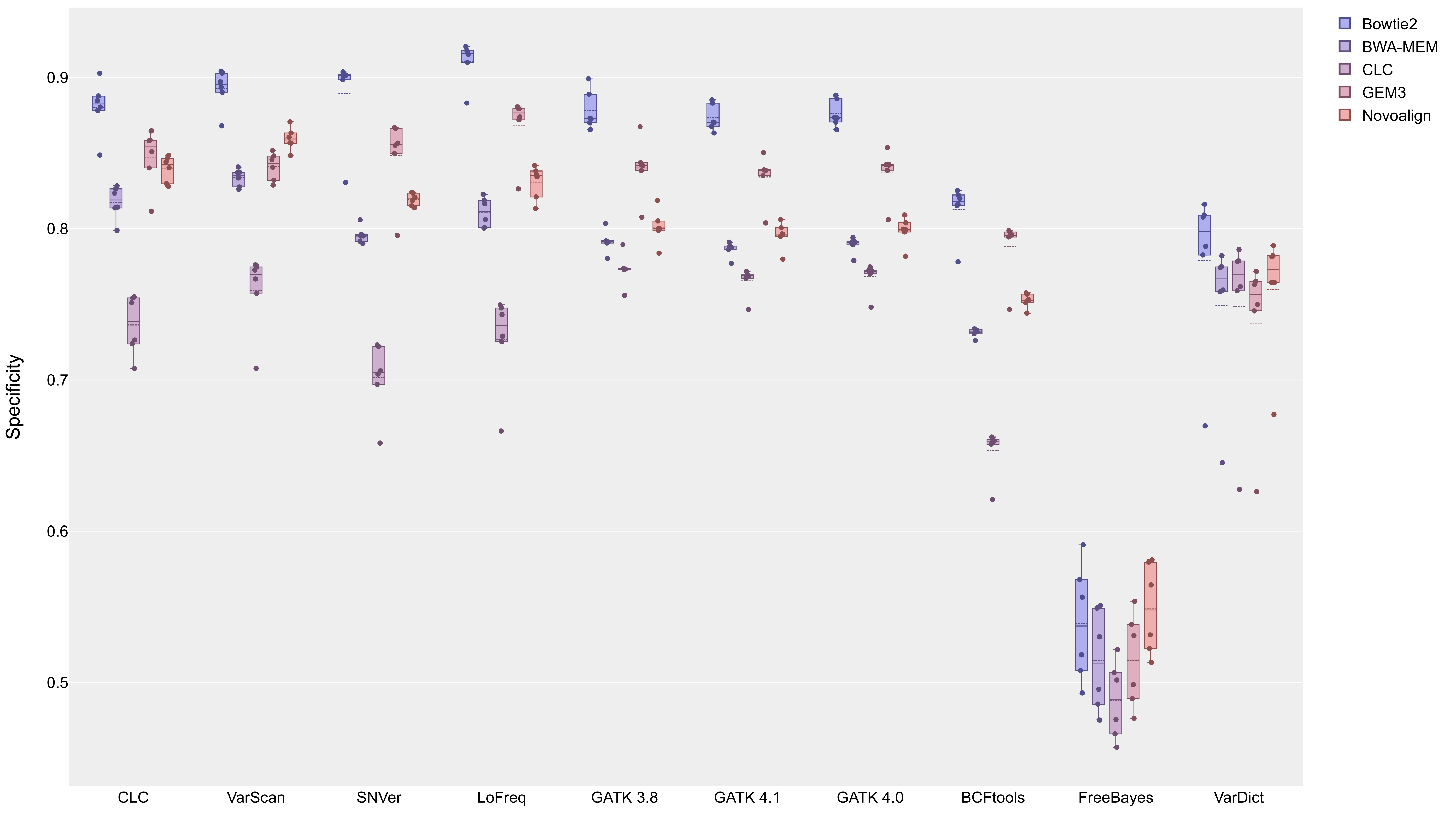

### Figure S4

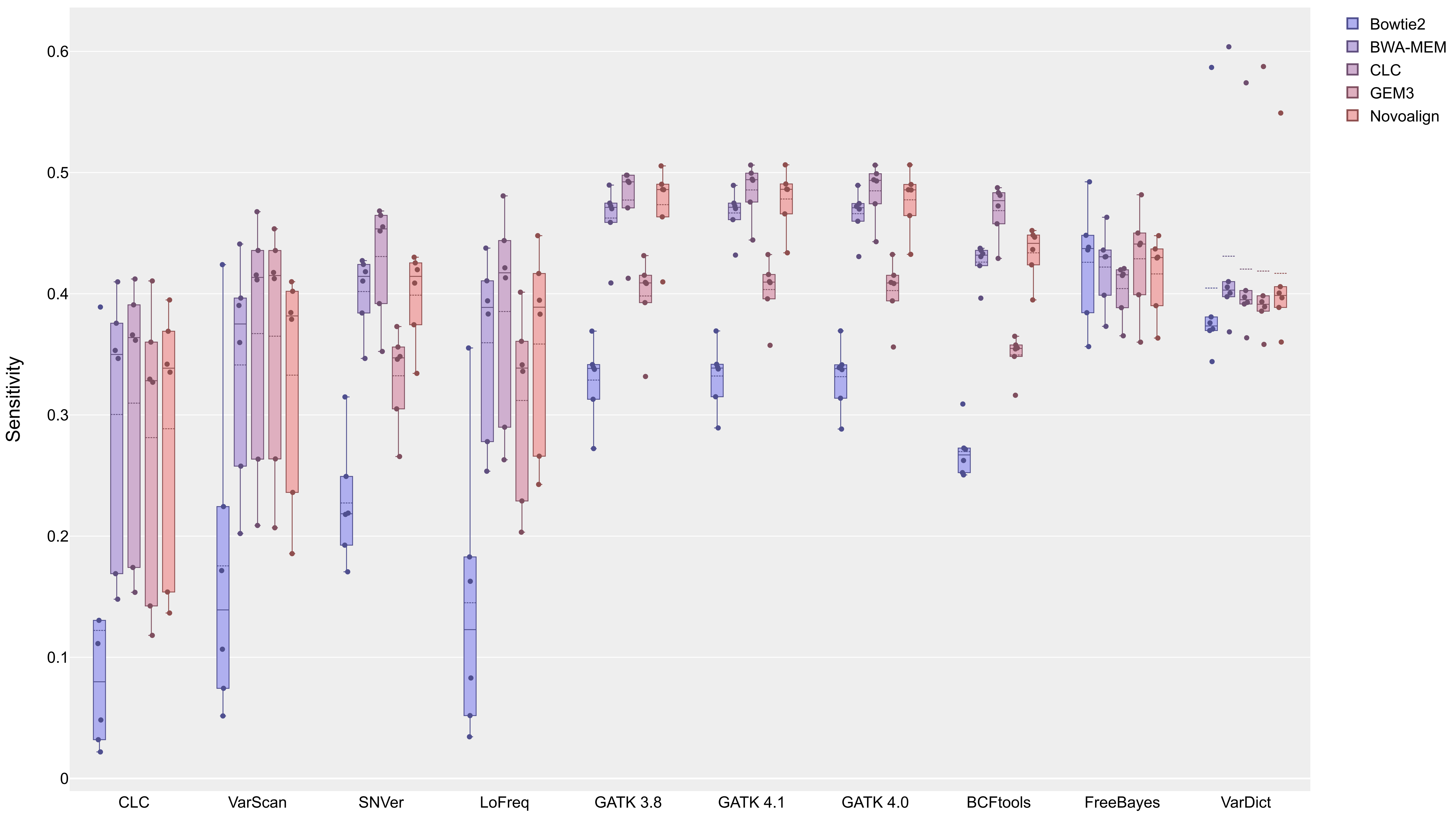

### Figure S5

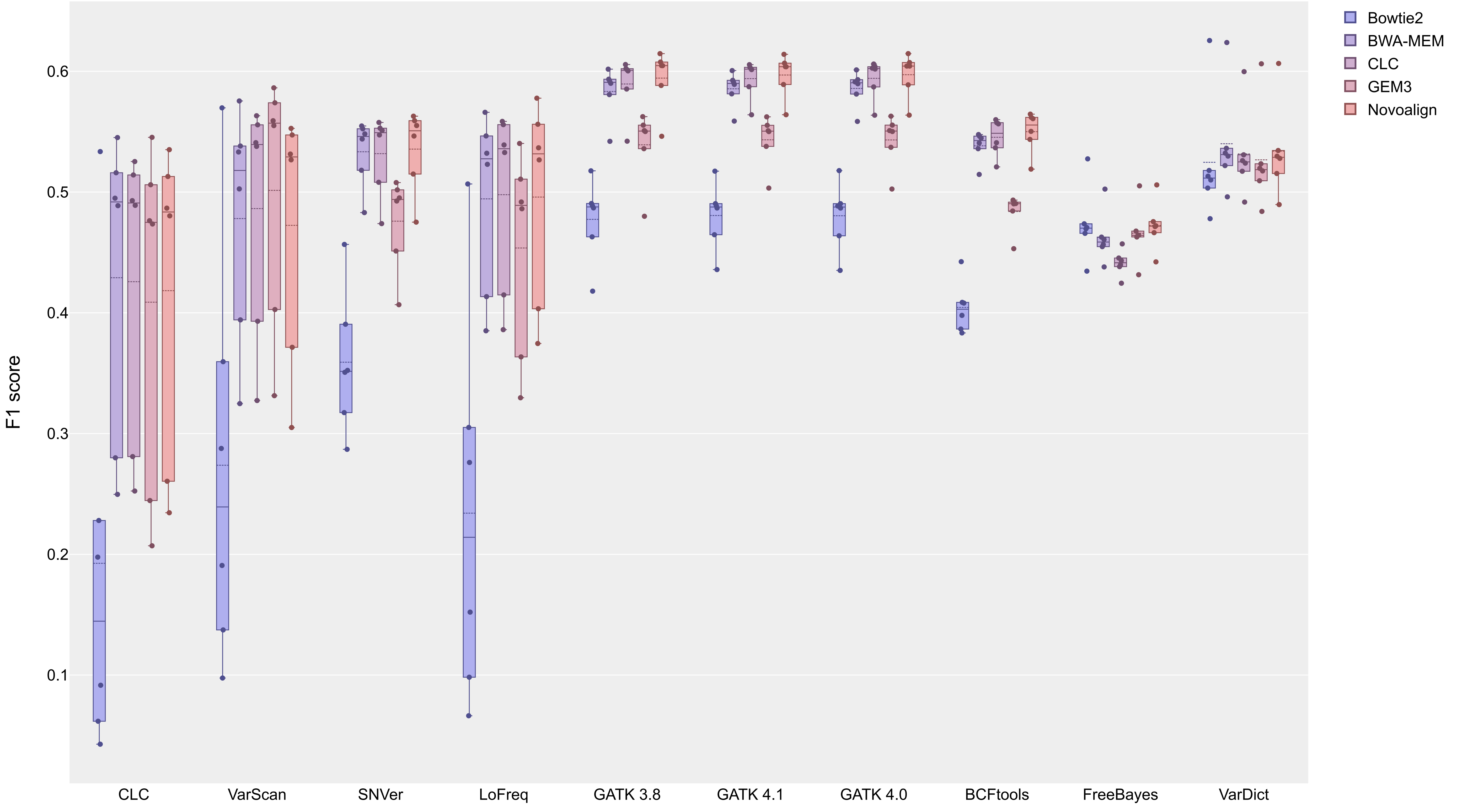

### Figure S6

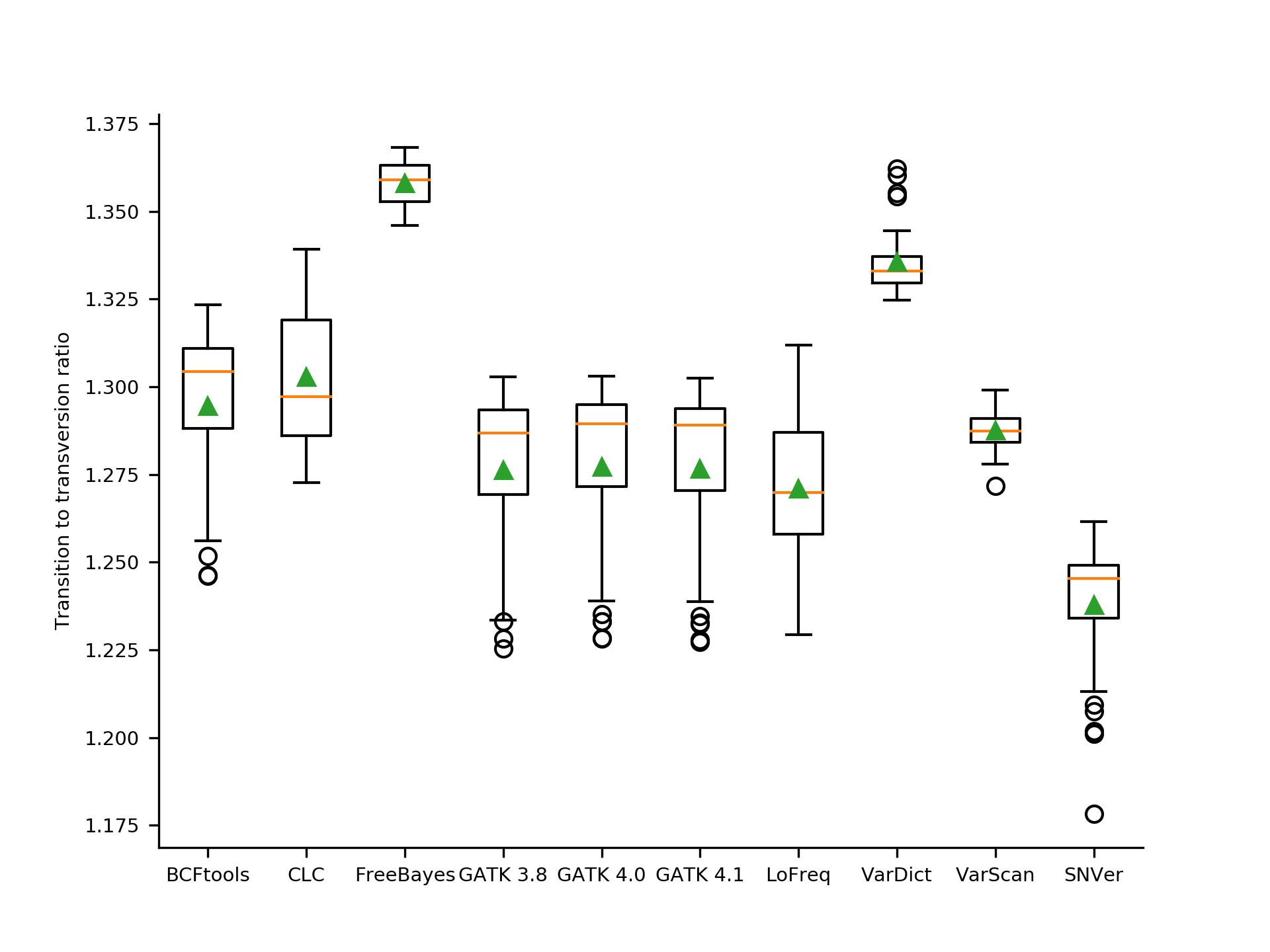
